# Exploring Stop Signal Reaction Time Over Two Sessions of the Anticipatory Response Inhibition Task

**DOI:** 10.1101/2022.07.21.500981

**Authors:** Alison Hall, Ned Jenkinson, Hayley J. MacDonald

## Abstract

Various behavioural tasks can measure the motor component of impulse control, response inhibition. Response inhibition encompasses the ability to cancel unwanted actions and is evaluated via stop signal reaction time (SSRT). The current study explored the effect of two sessions on SSRT within the anticipatory response inhibition task (ARIT) and how this compared to the stop signal task (SST). Forty-four participants completed two sessions of the ARIT and SST, 24 hours apart. SSRT and its constituent measures (Go trial reaction time, stop signal delay) were calculated. SSRT reflecting non-selective inhibition was consistent between sessions in both tasks (both p > .293). Reaction time and stop signal delay also remained stable across sessions in the ARIT (all p > .063), whereas in the SST, both reaction time (p = .013) and stop signal delay (p = .009) increased. Across the two sessions, SSRT reflecting partial inhibition improved (p < .001), which was underpinned by changes to reaction time (p < .001) and stop signal delay (p < .001). The maximal efficiency of non-selective inhibition remained stable across two sessions in the ARIT. Results of the SST confirmed that non-selective inhibition can however be affected by more than inhibitory network integrity when Go trial reaction times are not constrained in task design. Partial response inhibition measures changed across sessions, suggesting the sequential process captured by the SSRT occurred more quickly in session two. These findings highlight the absence/extent of inherent SSRT changes possible during multiple-session study designs e.g., pre/post, or active/sham comparisons.

## INTRODUCTION

Response inhibition (RI) is the motor component of inhibitory control and encompasses the ability to supress or cancel unwanted actions. There are various behavioural tasks used to objectively measure RI. One of the most popular is the stop signal paradigm which was first created by Vince (Vince, 1948), further developed into the stop signal task (SST) by Lappin & Eriksen (Lappin and Eriksen, 1966). Most recently the SST has been popularised by Logan and colleagues via the open-source STOP-IT software (Windows executable software for the stop-signal paradigm) which was first developed in 2008 (Verbruggen et al., 2008) and recently updated (Verbruggen et al., 2019). The SST involves Go trials where participants form a conditioned response to a Go signal, which is often a choice response. Participants respond to this Go signal as fast as possible with the press of a specific button. The SST also contains Stop trials making up approximately 25% of total trials. During Stop trials, a visual or auditory signal is presented after the Go stimulus and participants must inhibit their conditioned response. The task uses a staircase design to adapt this stop signal to the performance of the participant, narrowing in on a stop signal delay (SSD) where the participant successfully withholds their response on 50% of the stop trials. Logan and Cowan (Logan and Cowan, 1984) posited a horse-race model of RI to explain the behavioural outcome on each trial of the SST. The horse-race model suggests a race between the going process (initiated by the Go signal) and stopping process (initiated by the Stop signal) on a trial-by-trial basis. If the going process finishes first, then the response is executed, but if the stopping process finishes first, the responses is inhibited. Stop signal reaction time (SSRT) is the most widely utilised primary dependent measure for the SST as it is thought to indicate the latency of this stopping process/RI for an individual (Band et al., 2003; Aron and Poldrack, 2006; Li et al., 2008; Verbruggen and Logan, 2009; Verbruggen et al., 2013) .

Another method for investigating RI is via the anticipated response version of the SST. This version was developed by Slater-Hammel (Slater-Hammel, 1960) and follows the same horse race framework(Leunissen et al., 2017). This version of the SST constrains the Go response to an anticipated stationary target, to ensure response preparation takes place on both Go and Stop trials. When a Stop signal is presented before this anticipated target, participants must inhibit their Go response. Early versions of this anticipated response task, commonly named the anticipatory response inhibition task (ARIT), presented a clock face display and participants depressed a key to initiate a clockwise sweep dial revolution (Stinear and Byblow, 2004; Coxon et al., 2006). During Go trials, participants were required to release the key when the dial intercepted the target, 800ms after the start of the trial (Go response). Stop trials commenced in the same way, but the sweep dial stopped revolving before the target (Stop signal). Participants therefore had to inhibit their anticipated response when the Stop signal (the sweep hand stopping) was presented. More recent studies typically use a version of the ARIT involving one or two vertical bars which rise for 1000ms e.g. (Coxon et al., 2007, 2009; Zandbelt and Vink, 2010; MacDonald et al., 2012; Coxon et al., 2016; MacDonald et al., 2016; Gilbert et al., 2019; He et al., 2019). This version of the ARIT has been increasing in popularity and is now available open-source (He et al., 2021) In the bimanual version, participants are required to release two depressed keys to intercept two bars with the target line at 800ms on Go trials. Participants then inhibit this bimanual lift response when the bars do not reach the target on Stop Both trials, with the latency of the non-selective stopping process reflected in the SSRT. During the more challenging Partial Stop trials, participants are required to keep only one key depressed when the corresponding bar does not reach the target and release the alternative key at the target line. The response of the continuing hand is invariably delayed, which is termed the stopping interference effect (Aron and Verbruggen, 2008; Ko and Miller, 2011; Wadsley et al., 2019). SSRTs are also calculated on these trials but are thought to reflect a more complex series of neural processes triggered by the stop signal; a sequential non-selective stop, uncouple and reprogram, then selective go process (Coxon et al., 2009; MacDonald et al., 2012; MacDonald et al., 2014; Wadsley et al., 2019; MacDonald et al., 2021).

Research conducted using both the ARIT and SST has revealed the neural mechanisms underlying inhibitory control, specifically the role of basal ganglia pathways. Execution of the motor response in Go trials activates fronto-striato-pallidal regions as part of the direct basal ganglia pathway, which then leads to an increase in thalamocortical drive to the motor cortex. Whereas inhibition on Stop trials engages a right lateralized network that includes the indirect (suppression of action) or hyperdirect (cancellation of action) pathway, that inhibits output from the motor cortex. This inhibitory network includes the subthalamic nucleus (STN), globus pallidus pars interna (GPi) and externa (Gpe) (indirect pathway), right inferior frontal gyrus (IFG) and pre-supplementary motor area (SMA) (Aron et al., 2003; Aron and Poldrack, 2006; Li et al., 2008; Coxon et al., 2009; Ray et al., 2012; Dunovan et al., 2015; Allen et al., 2018; Chen et al., 2020; Maizey et al., 2020). The STN, once activated via the indirect or hyperdirect pathway, plays an important role in supressing thalamocortical output by blocking the direct pathway (Aron and Poldrack, 2006; Li et al., 2008; Zandbelt and Vink, 2010; Dunovan et al., 2015)(. The integrity of these basal ganglia pathways is thought to be reflected in measures derived from RI tasks.

There is substantial literature investigating single session measures of RI using the ARIT and SST. The assumption is that RI, indexed via a SSRT, is an inherent ability which is specific to each individual and purely reflects the integrity of their inhibitory networks. Therefore, RI is not expected to change within a young healthy individual and SSRT should be consistent across multiple sessions. A handful of previous studies have tested this assumption by investigating the effect of multiple sessions on RI in the SST. Two of these studies reported no improvement in SSRT for non-selective RI following 9 and 2 sessions, respectively (Enge et al., 2014; Chowdhury et al., 2020). While Chowdhury et al. found no behavioural improvement in stopping efficacy with multiple sessions, Enge et al. counterintuitively reported an increase in SSRT (i.e., a decrease in performance) over the course of multiple sessions. Enge and colleagues attributed this to participants progressively focusing more on fast responses in Go trials at the expense of accurate cancellation on Stop trials. This interpretation suggests strategizing during the task might be able to affect the SSRT measurement over multiple sessions. Conversely, another study using the SST for non-selective RI did find an improvement in SSRT throughout 10 sessions, where SSRT decreased with each session (Berkman et al., 2014). SSRT for partial RI on the SST has shown a similar pattern of decreasing across two sessions, which trended towards significance (Xu et al., 2014). To our knowledge, no study has specifically tested SSRT across multiple sessions of the ARIT. Coxon and colleagues (Coxon et al., 2016)reported behavioural results on the ARIT pre and post neuroimaging. Although SSRT was not reported as it was not a primary outcome measure, they did show that Go trial RT remained stable across the two behavioural sessions and only RT variability (1SD of response distribution) significantly decreased. To ensure we are correctly interpreting SSRT as an inherent measure of inhibitory network integrity, the consistency of SSRT across multiple sessions needs to be further explored.

The aim of the current study was therefore to assess if and how the SSRT measurement changed over two sessions for the ARIT, and how this compared with the SST. It was hypothesised that SSRT would not change between sessions on Stop Both trials of the ARIT as there would be no possible change in Go trial reaction times used to calculate this measure due to constraining responses to a stationary target. It was suspected that a strategy focusing on accurate stopping at the expense of fast reaction times on Go trials of the SST (i.e. the opposite strategy to participants in the Enge et al. 2014 study) would be able to cause an improvement in inhibitory control, as seen previously (Berkman et al., 2014). Therefore, the second hypothesis was that SSRT in the SST would decrease from session one to session two. Due to the increased challenge at both a behavioural and neural level on Partial Stop trials in the ARIT, becoming better at fulfilling the trial requirements in session two might affect performance. Therefore, the third hypothesis was that SSRT would decrease from session one to session two for Partial Stop trials of the ARIT.

## MATERIALS AND METHODS

### Participants

Forty-four healthy participants were recruited into the current study, all over the age of 18 years. The University of Birmingham Ethics Committee approved this research and written informed consent was obtained from each participant.

### Procedure

Participants attended two identical sessions in the laboratory, 24 hours apart. Participants were seated ~1m away from a computer screen and keyboard, where they completed two behavioural tasks: the anticipatory response inhibition task (ARIT) and the stop signal task (SST). The order of behavioural tasks was counterbalanced.

### Anticipatory Response Inhibition Task (ARIT)

The ARIT was displayed using custom code written in MATLAB (version R2016a, MathWorks). Participants completed two practise blocks, each containing 30 Go trials, followed by 6 experimental blocks of 30 trials. In total the experimental trials consisted of 120 Go trials and 60 Stop trials in a pseudo-randomised order.

Participants were initially presented with a grey screen containing two white vertical rectangles and a stationary horizontal black target line 4/5 of the way up the rectangles (Figure 1). All trials required participants to use their left and right index fingers to depress the ‘z’ and ‘?/’ key, respectively. Once both keys were depressed, a black bar started rising within each of the white rectangles after a variable delay. The left black bar was controlled with the ‘z’ key, and the right black bar with the ‘? /’ key. Both bars rose at equal rates, intercepting the target (horizontal black line) at 800ms and filling the entire white rectangle at 1000ms, unless the keys were released which ceased the bars rising.

**Figure 1.**
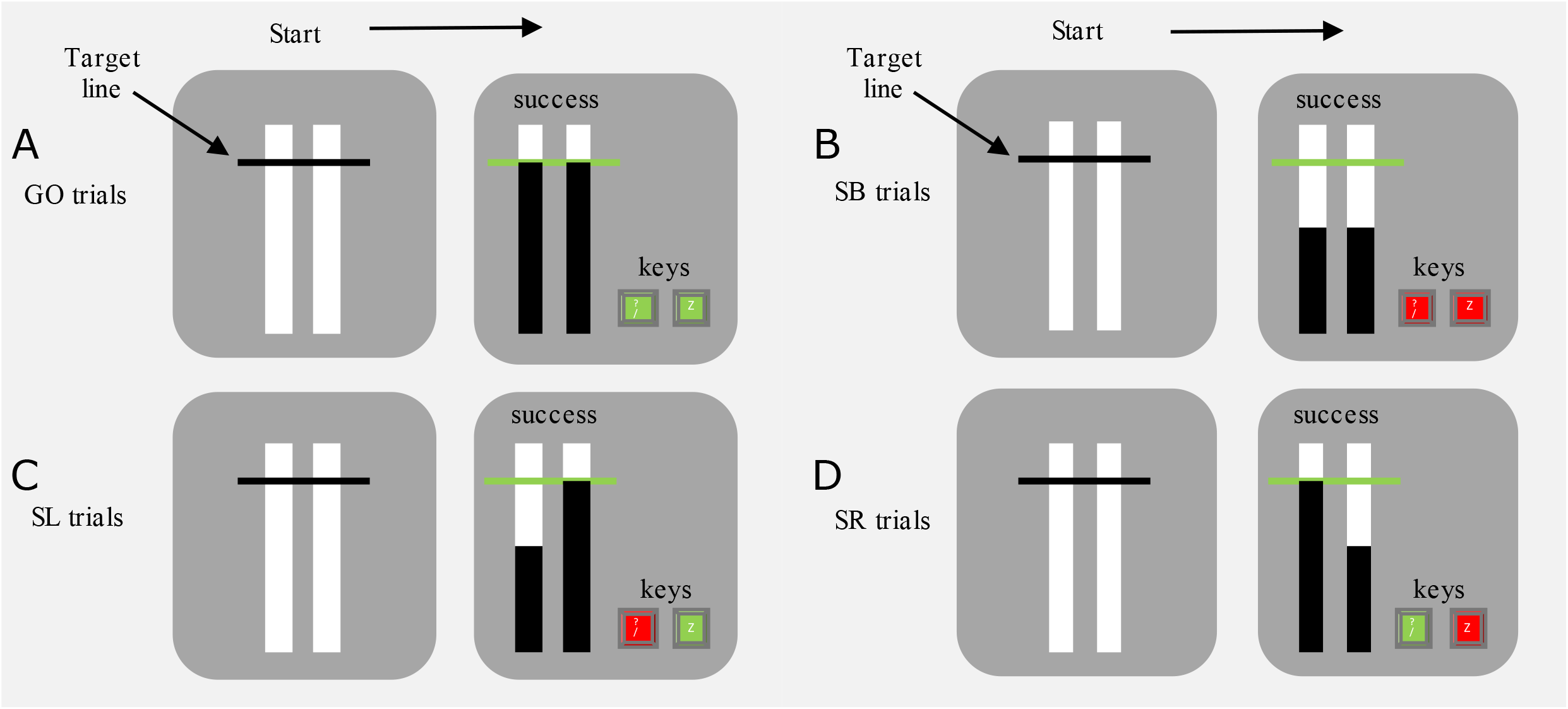
Visual display of (**A**) GO, (**B**) SB (stop both), (**C**) SL (stop left) and (**D**) SR (stop right) trials in the ARIT. Green keys represent successful release at the target and red keys represent successfully keeping the key depressed. On successful Go trials, both keys are released at the target line. On successful SB trials, both keys are held down. On successful SL trials, the right key is lifted and the left key is held down. On successful SR trials, the left key is lifted and the right key is held down.

#### Go trials

Participants started each trial by pressing and holding down both response keys to initiate the rising of the bars. They were then required to release their fingers from both keys to intercept both bars with the target (successful releases were within 30ms of target, Figure 1A). Participants received visual feedback at the end of each trial which was shown above the white rectangles. ‘Success’ was displayed following a successful release of the keys and ‘missed’ followed an incorrect release (not within 30ms of target).

#### Stop trials

There were three types of Stop trials (20 each) that required participants to keep the key(s) depressed if the rising bar(s) never reached the target. Stop both (SB) trials required participants to keep both keys depressed when both bars automatically stopped rising before reaching the target (non-selective/complete RI) (Figure 1B). Partial Stop trials comprised of stop left-go right (SL) and stop right-go left (SR) trials (Figure 1C & D), which required participants to keep pressing the key corresponding to the bar that stopped, whilst still releasing their finger from the alternative key when the bar arrived at the target (partial RI). For every stop version, the bar initially stopped 550ms into the trial (stop signal delay, SSD). A staircase algorithm with increments of 50ms was then used to generate a 50% success rate for each Stop trial version. Following a successful Stop trial the SSD would increase by 50ms for the subsequent Stop trial, whereas the SSD would decrease by 50ms following an unsuccessful Stop trial. Participants received feedback following each trial which was displayed above the white rectangles. Following successful trials, ‘success’ was displayed, and ‘unsuccessful stop’ was presented following an unsuccessful trial where participants did not inhibit their response. Moreover, on Partial Stop trials, if the participants completed a successful stop on the required side and released the alternative key outside 30ms of the target, then ‘successful stop, but missed target’ appeared (these results were classed as a successful stop in the analyses as this delayed response - i.e. stopping interference effect - was expected).

### Stop Signal Task (SST)

The SST was carried out using STOP-IT software (Verbruggen et al., 2008). Participants completed a practise block of 48 Go trials and 16 Stop trials. They then completed 128 experimental trials, divided into 2 blocks. In total these trials consisted of 96 Go trials and 32 Stop trials in a pseudo-randomised order.

#### Go trials

The start of a trial was indicated by a white ‘+’ fixation cue (approximately 1cm across) displayed for 250ms in the centre of the computer screen with a black background. This cue was followed by the presentation of the Go stimulus. The Go stimulus was either a white square or circle (approximately 1cm in length) which required participants to respond with their left or right index finger as fast as possible using the ‘z’ or ‘? /’ key, respectively (Figure 2A). Participants had up to 1250ms to respond to the Go stimulus once it was presented, before the trial ended and the stimulus disappeared. A blank screen would be displayed for 750ms before the start of the subsequent trial.

**Figure 2.**
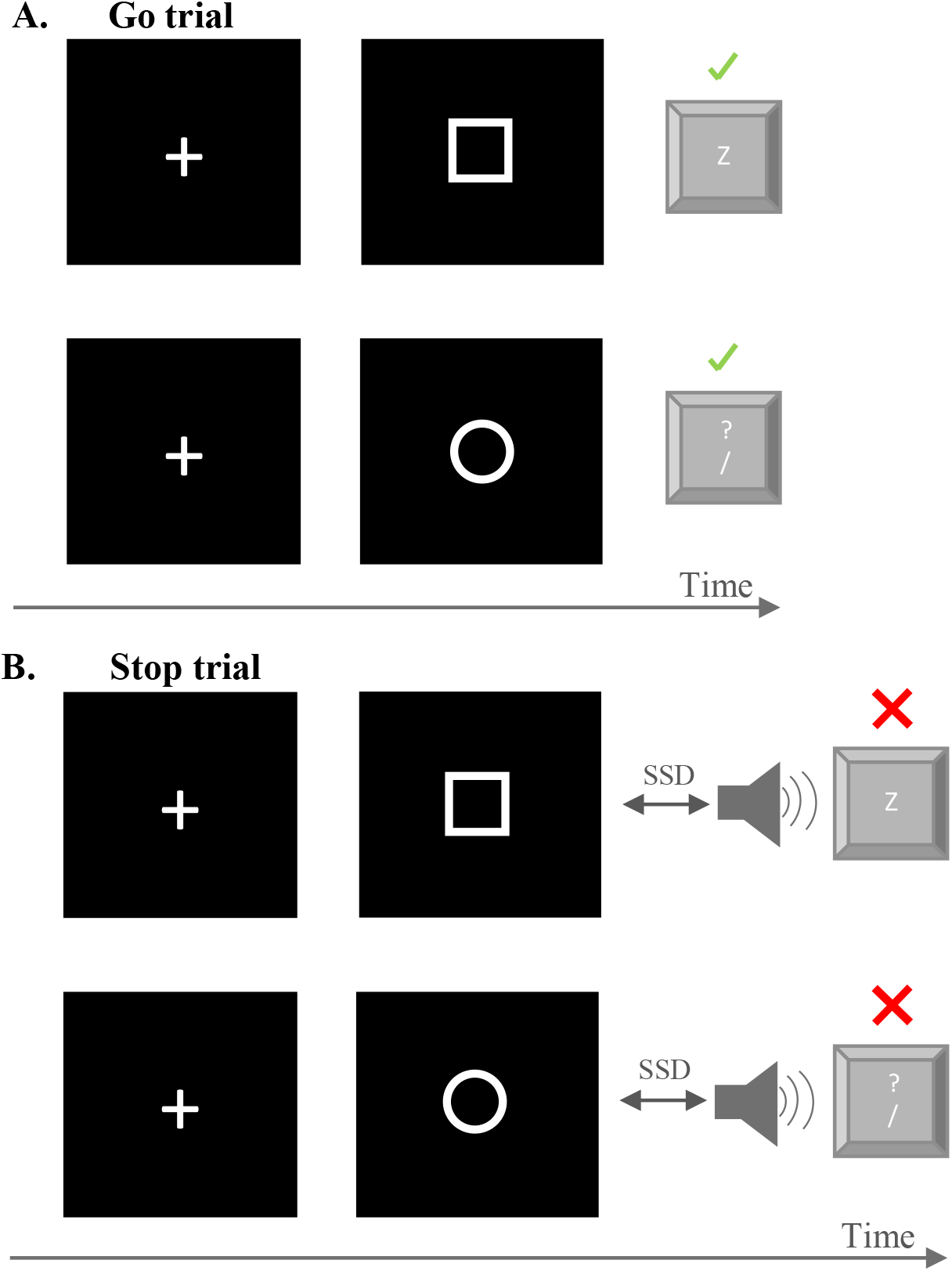
Visual representation of the Stop Signal Task. Green tick marks represent a successful Go trial (**A**) where the participant presses the correct key corresponding to the symbol. Red crosses represent a successful Stop trial (**B**) where participants refrain from pressing the key following the stop signal.

#### Stop trials

Stop trials also commenced with the same ‘+’ on the computer screen indicating the start of the trial, followed by a blank screen and subsequently the presentation of the Go stimulus. Shortly after the Go stimulus, a Stop stimulus was presented, which was a short audio tone lasting 75ms (750hz). Participants were instructed to inhibit their response and not depress the designated key (Figure 2B) if they heard the Stop tone (non-selective/complete RI). The time between the Go stimulus and the Stop stimulus represented the stop signal delay (SSD). The SSD was initially set at 300ms after the Go stimulus and a staircase algorithm was used to generate a 50% success rate, where the SSD would increase by 50ms on the subsequent Stop trial (regardless of whether the Go response was with the left or right hand) if the participant successfully inhibited their response, but the SSD would decrease by 50ms if the participant was unsuccessful (i.e. responded following the stop stimulus). The trial would end after 1250ms and the Go stimulus would disappear. There was an equal number of Go and Stop trials for each hand response. Mean reaction time and percentage of correct stops were provided as feedback at the end of each block and were displayed for 10 seconds.

### Dependent measures

#### Anticipatory response inhibition task

Average reaction time (RT), reported in milliseconds relative to the start of the trial, was calculated for successful Go, SL (stop left-go right) and SR (stop right-go left) trials after removing outliers (±3SD, MacDonald, Stinear & Byblow, 2012). SSD (staircased to 50% success) and stop trial accuracy (% success) was calculated for SB SL, SR trials. SSRT was the primary dependent measure for the ARIT and calculated for each Stop trial version using the integration method (SSRT = nth Go trial RT (i.e. number of Go trials x probability of responding on Stop trials) – SSD) (Verbruggen et al., 2013).

#### Stop signal task

The RT, reported in milliseconds, was measured between the onset of the Go stimulus and the key response. Average RT across Go stimuli (outliers of ±3SD were removed for consistency between tasks), SSD (staircased to 50% success) and stop trial accuracy (% success) on Stop trials were calculated for each participant. SSRT was the primary dependent measure for the SST and was also calculated using the integration method.

### Statistical Analysis

MATLAB (Version R2020a, MathWorks) and SPSS statistics (Version 27) were used to complete all statistical analyses. To investigate the effect of session on non-selective inhibitory control, a direct comparison was made between Stop Both trials in the ARIT and the Stop trials in the SST due to the similar requirement for complete RI in both conditions. A 2 Session (First, Second) x 2 Task (ARIT SB, SST) repeated measures analysis of variance (RM ANOVA) was run on SSRT and SSD. A similar Session x Task (ARIT, SST) RM

ANOVA was run on average Go trial RT, which is a key measure used to calculate SSRT. To investigate the effect of session on partial RI in Partial Stop trials of the ARIT, a 2 Session x 2 Partial Stop Type (SL, SR) RM ANOVA was run on SSRT, SSD and RT from these trials. Post hoc paired t-tests were used to investigate any significant main effects and interactions. One sample t-tests were used to compare the percentage of stop trial success (stop trial accuracy) in the ARIT (SB, SL, SR) and SST to the 50% staircasing target, and paired t-tests were used to assess the differences in stop trial accuracy from session 1 to session 2 for all trial types.

To investigate the generalizability of SSRT across tasks, linear regressions tested for a correlation between SSRTs calculated in the SB trials of the ARIT and the SST Stop trials, for each session. Fisher z transformations identified any significant differences between the correlations. Values are reported as means ± standard error (SE) unless otherwise stated. Statistical significance was determined by α ≤ 0.05 and partial eta squared effect sizes are reported. Data which violated the assumption of sphericity are reported with Greenhouse-Geisser corrected p values.

## RESULTS

### ARIT Stop Both and SST

Eight participants were excluded from the main analysis due to SSRTs in the SST being below 100ms which is not feasible for reactive recruitment of the inhibitory network (advised by (Congdon et al., 2012; Verbruggen et al., 2019)). Data from the remaining 36 participants were used to test the first two hypotheses (mean age: 20 ± 0.94 years, range 18-22 years, 12 males).

#### Stop signal reaction time (SSRT)

SSRT reflecting non-selective inhibitory control appeared to be consistent between the two sessions when measured using either the SST or ARIT. There was no main effect of Task (F_1,35_ = 0.04, p = .840, ηp^2^ = .001), Session (F_1,35_ = 1.14, p = .293, ηp^2^ = .032) or Task x Session interaction (F_1,35_ = 1.82, p = .186, ηp^2^ = .049) (Figure 3A). As hypothesised for Stop Both trials of the ARIT, SSRT did not change between session one (226 ± 10ms) and session two (226 ± 77ms). Contrary to our second hypothesis, the decrease in SSRT from session one (236 ± 18ms) to session two (212 ± 11ms) in the SST was not significant. While an individual’s SSRT for non-selective RI was comparable between tasks initially, this relationship was not sustained into the second session. There was a significant positive correlation between SSRT on the ARIT and SST in session one (r = 0.45, p = .006) but this disappeared in session two (r = −0.07, p = .702) (Figure 3B). Fisher z transformation confirmed a significant difference between the two correlations (z = 2.22, p =.013).

**Figure 3.**
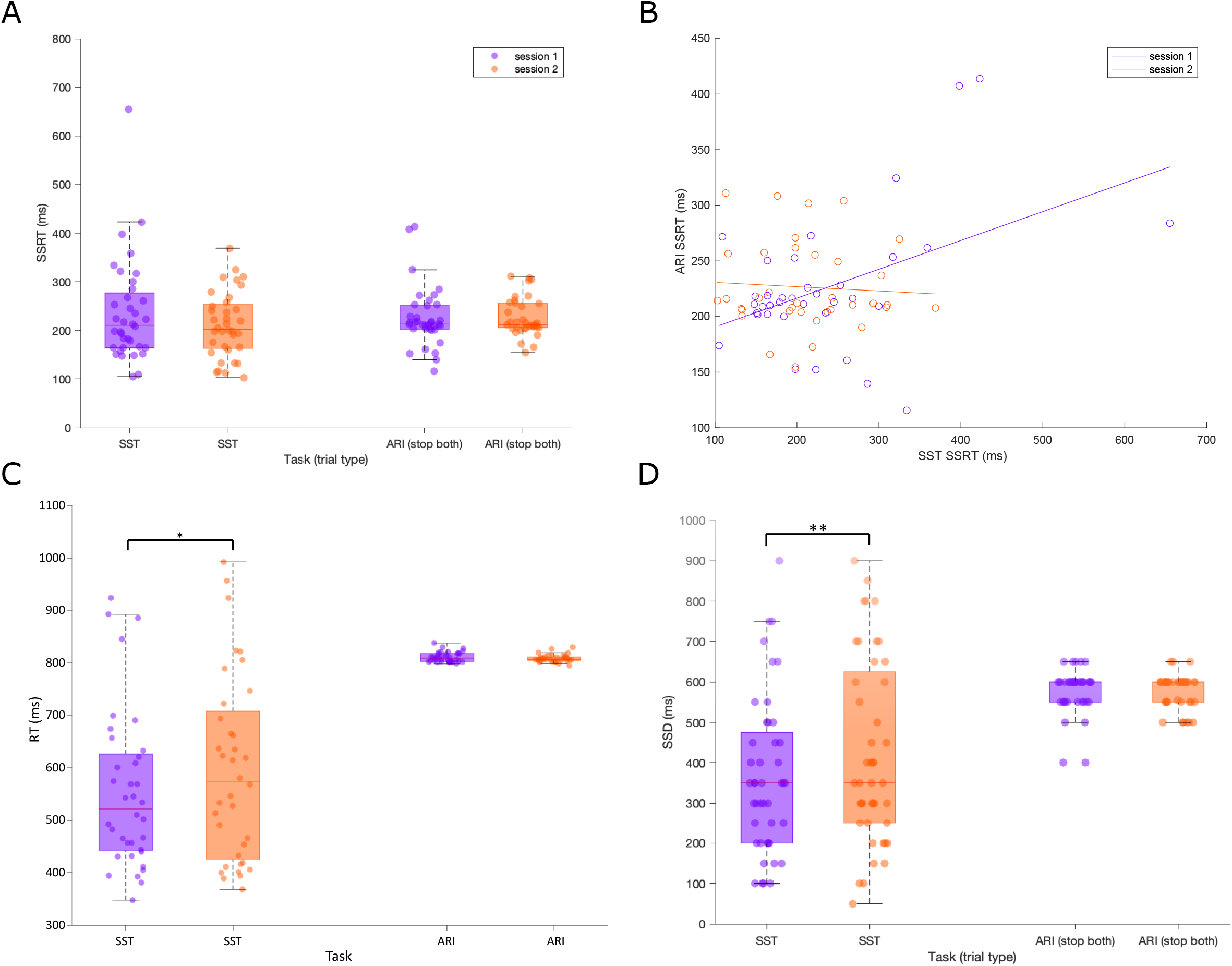
Mean SSRT **(A)**, RT **(C)** and SSD **(D)** for the ARIT SB and SST in sessions 1 and 2, reported in milliseconds (ms). Shaded box plots represent the interquartile range (IQR) (75^th^ percentile (Q3) – 25^th^ percentile (Q1)). Red horizontal line represents the median. The vertical dashed lines represent the non-outlier minimum (Q1 – 1.5 x IQR) and maximum (Q3 + 1.5 x IQR). Data circles represent individual participant results. **(B)** Linear correlation between SSRT for the SST and ARIT SB in session 1 and session 2. Data circles represent individual participant SSRT. * p < .05, ** p < .01.

#### Reaction time (RT)

Reaction times on Go trials remained consistent over two sessions in the ARIT but were significantly delayed in session two of the SST. There was a main effect of Task (F_1,35_ = 80.8, p < .001, ηp^2^ = .698), Session (F_1,35_ = 6.00, p = .019, ηp^2^ = .146) and Task x Session interaction (F_1,35_ = 7.77, p = .009, ηp^2^ = .182). Post hoc analysis revealed the mean decrease of 2ms from session one (811 ± 2ms) to session two (809 ± 1ms) in the ARIT was not significant (p = .063). However, the increase of 41ms from session one (555 ± 25ms) to session two (596 ± 29ms) in the SST was significant (p = .013; Figure 3C).

#### Stop signal delay (SSD)

Mirroring the RT results, the SSD was longer in session two of the SST but remained consistent for the ARIT. There was a main effect of Task (F_1,35_ = 56.1, p < .001, ηp^2^ = .616), Session (F_1,35_ = 5.02, p = .032, ηp^2^ = .125) and Task x Session interaction (F_1,35_ = 6.94, p = .012, ηp^2^ = .165). Again, post hoc analysis revealed no significant changes in SSD for the ARIT (session one = 582 ± 9ms, session two = 583 ± 7ms; p = .884), but a significant increase in SSD for the SST (session one = 329 ± 30ms, session two = 375 ± 35ms; p = .009; Figure 3D).

#### Stop trial accuracy

There were no differences in stop trial accuracy between session one and session two in either task. In the ARIT there was a non-significant increase of 0.14 ± 0.61% between session one (51.5 ± 0.56%) and session two (51.7 ± 0.40%) (t_1,35_ = 0.23, p = .822). Additionally, in the SST there was a non-significant increase of 1.61 ± 1.58% between session one (49.9 ± 2.46%) and session two (51.6 ± 0.81%) (t_1,35_ = 1.02, p = .314). When comparing mean stop trial accuracy in both sessions to the 50% staircasing target, only the ARIT task displayed percentages significantly greater than 50% (both p < .01).

### ARIT Partial Stop Trials (Stop Left and Stop Right)

Data from the full 44 participants were included in the following analyses (mean age: 20 ± 1 years, range: 18-22 years, 15 males).

#### Stop signal reaction time

SSRT reflecting partial RI improved over the course of two sessions. There was a main effect of Session (F_1,43_ = 31.4, p = <.001, ηp^2^ = .422) with a decrease of 98 ± 17ms in SSRT from session one (451 ± 21ms) to session two (353 ± 19ms; Figure 4A). There was no main effect of Stop type (F_1,43_ = 1.76, p = .192, ηp^2^ = .039) or Stop type x Session interaction (F_1,43_ = .128, p = .722, ηp^2^ = .003).

**Figure 4.**
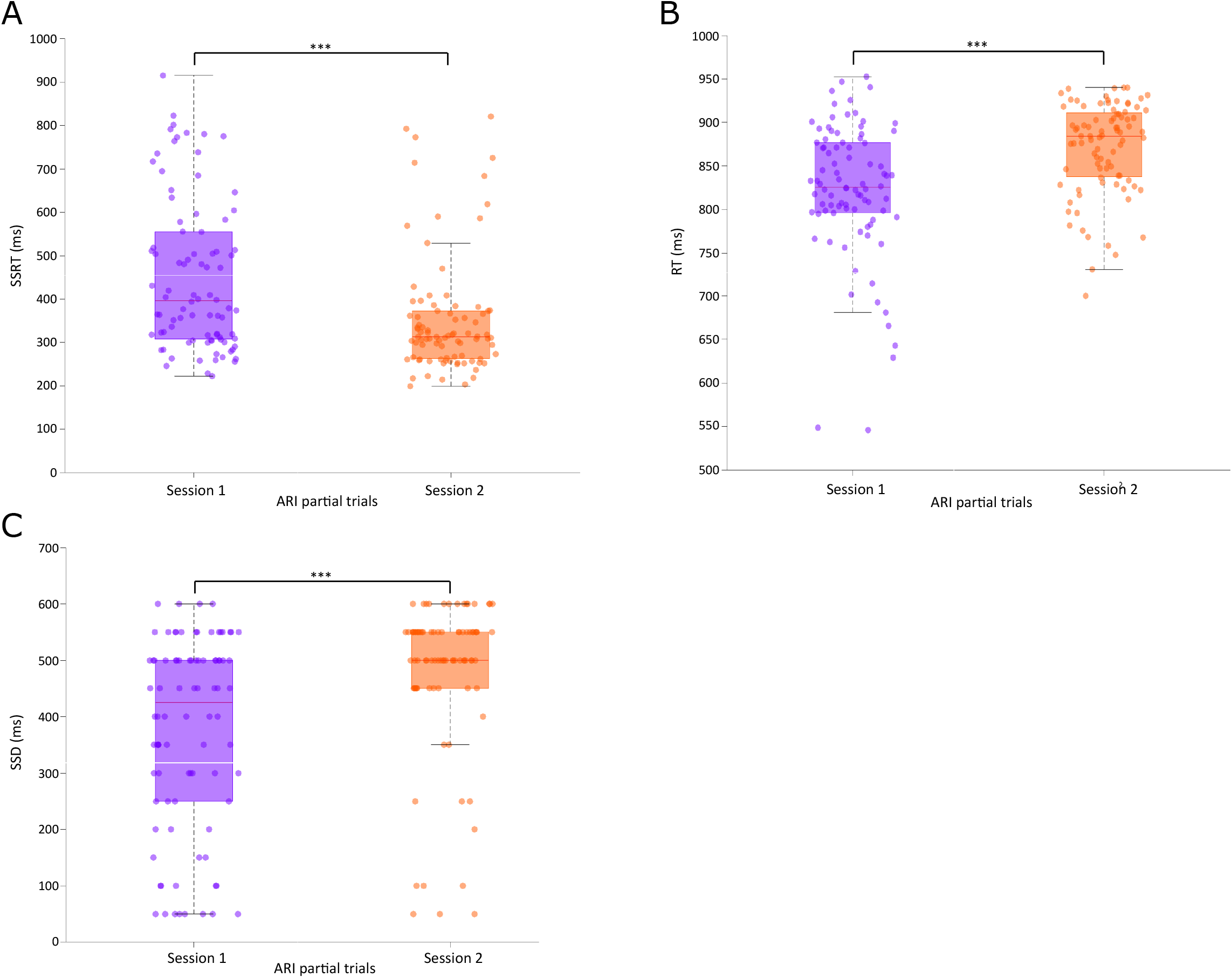
Mean SSRT **(A)**, RT **(B)** and SSD **(C)** for partial response inhibition on the ARIT (partial trials collapsed across side) in sessions 1 and 2, reported in milliseconds (ms). Shaded box plots represent the interquartile range (IQR) (75^th^ percentile (Q3) – 25^th^ percentile (Q1)). Red horizontal line represents the median. The vertical dashed lines represent the non-outlier minimum (Q1 – 1.5 x IQR) and maximum (Q3 + 1.5 x IQR). Data circles represent individual participant results. *** p < .001.

#### Reaction time

Regardless of which side was still responding on Partial Stop trials, there was a larger delay in RT relative to the target in session two compared to session one. There was a main effect of Session (F_1,43_ = 36.1, p = <.001, ηp^2^ = .456) from a significant increase of 48 ± 8ms in RT from session one (822 ± 10ms) to session two (870 ± 7ms; Figure 4B). There was no main effect of Stop type (F_1,43_ = .825, p = .369, ηp^2^ = .019) or Stop type x Session interaction (F_1,43_ = 1.59, p = .214, ηp^2^ = .036).

#### Stop signal delay

In a similar pattern as RTs, SSD increased over the two sessions during both types of partial RI. There was no main effect of Stop type (F_1,43_ = 1.70, p = .199, ηp^2^ = .038) or Stop type x Session interaction (F_1,43_ = .462, p = .501, ηp^2^ = .011) but there was a main effect of Session (F_1,43_ = 37.5, p = <.001, ηp^2^ = .466). There was a significant increase of 106 ± 17ms in SSD from session one (374 ± 21ms) to session two (480 ± 19ms; Figure 4C).

#### Stop trial accuracy

The stop trial accuracy improved over the two sessions during both types of partial RI. For SL, there was an increase of 4.39 ± 1.65% from session one (42.0 ± 1.90±) to session two (46.4 ± 1.56%) (t_1,43_ = 2.66, p = .011). For SR, there was an increase of 5.18 ± 1.74% from session one (40.6 ± 1.56%) to session two (45.8 ± 1.20%) (t_1,43_ = 2.98, p = .005). Both types of partial RI displayed a mean stop trial accuracy significantly lower than 50% in both sessions (all p < .027).

## DISCUSSION

The current study constitutes the first step towards exploring the consistency of SSRT across multiple sessions of the ARIT. Performing two experimental sessions of the ARIT had distinct effects on SSRT measured during complete versus partial RI. As hypothesised, there was no change over the two sessions for SSRT reflecting complete RI. This stability supports the idea that SSRT measuring non-selective inhibitory control on this task reflects the inherent ability of an individual to inhibit a response and is therefore not expected to change. The consistency of this measure was underpinned by no change in SSD for non-selective Stop trials as well as no change in RT on Go trials in this task. This finding was in contrast to inhibitory control on the SST which was associated with a longer SSD and delayed RT on Go trials in session two. However, contrary to our hypothesis, the comparable increases in both SSD and Go RT resulted in no significant change to SSRT overall across sessions of the SST. Conversely, SSRT reflecting partial RI on the ARIT decreased from session one to session two. This decrease was as predicted and observed because participants became better at fulfilling the demands on Partial Stop trials in session two, reflected by a longer SSD. Overall, these findings have implications for i) the extent of potential within-individual changes to SSRT during multiple-session study designs, and ii) how SSRT might be interpreted for complete versus partial RI.

Stop signal reaction time to a non-selective stop signal during the ARIT did not change across sessions. Importantly, the two variables used to calculate SSRT also remained constant. As such, the staircase algorithm employed successfully converged on a stop signal presentation time in the first session which reflected maximal efficiency of the inhibitory process. The prepotent anticipated response was also unaffected by session, replicating previous findings (Coxon et al., 2016), most likely from being constrained by the task design (Leunissen et al., 2017). The current findings suggest that SSRT calculated from Stop Both trials of the ARIT is indeed a valid measure of non-selective inhibition network activity. This measure would appear to fit Congdon and colleagues’ (2012) definition of SSRT as a “heritable measure of interindividual variation in brain function”. However, this cannot be confirmed by our study alone, and future studies should extend the number of sessions to ensure non-selective SSRT remains consistent. This is especially pertinent as the study by Berkman and colleagues (2014), despite constraining go responses on the SST, observed an improvement in RI throughout 10 sessions, as well as a proactive shift in the pattern of neural activation in RI networks. Therefore, despite the apparent robustness of movement execution and non-selective inhibition on the ARIT, future studies with a greater number of sessions and measures of neural activation are required to substantiate these findings.

Non-selective inhibitory control on the SST also appeared consistent between sessions. However, the consistency of the SSRT was belied by changes to SSD and Go RT across session. There was evidence of proactive slowing from session one to session two, reflected in the delayed going response. This slowing is purported to be an example of proactive motor RI (Schachar et al., 2004; Verbruggen et al., 2008; Verbruggen et al., 2013; Brevers et al., 2020)and could be attributed to participants focusing on successful stopping at the expense of fast responses on Go trials (i.e. the opposite strategy to participants in Enge et al. 2014). A strategy to prioritise stopping performance would also explain the longer SSD in session two, which indicates participants had improved at responding to the stop signal.

Importantly, employing such a strategy invalidates the independence assumption of the race model (Verbruggen et al., 2019) and might explain why non-selective inhibitory performance was no longer related between tasks in session two. Specific versions of the SST constrain responses to prevent such proactive slowing (Berkman et al., 2014; Chowdhury et al., 2020) and enable a more reliable interpretation of SSRT measures. Overall, when interpreting measures of inhibitory control on the SST, our results highlight the value in examining variables that constitute the SSRT despite no apparent change to SSRT itself, and that changes in these variables might suggest SSRT is able to be affected by more than purely inhibitory network integrity.

The SSRT measure needs to be interpreted differently for complete versus partial RI. This difference is not necessarily surprising as SSRT in Partial Stop trials is more than a measure of pure (or global; i.e stop everything) inhibitory network activity. The stop cue on these trials triggers a sequential non-selective stop, response uncouple, reprogram, then selective go process(Coxon et al., 2007; MacDonald et al., 2012; MacDonald et al., 2014; Cowie et al., 2016; Wadsley et al., 2019; MacDonald et al., 2021) which leads to the delayed RT. Partial RI SSRT therefore reflects a complex series of neural processes which are triggered by the partial stop cue and involve interactions between facilitatory and inhibitory prefrontal-basal ganglia networks (Coxon et al., 2009; Coxon et al., 2012). Of note, the fact that participants still make a unimanual response on Partial Stop trials means both components used to calculate SSRT (SSD, RT) can be measured within the same trial type. This is in contrast to SSRTs for Stop Both trials or Stop trials in the SST, which use RTs from Go trials as there is necessarily no overt response on successful Stop trials. In this way, SSRTs for partial RI are not comparable to SSRT from non-selective inhibition trials of the ARIT. The direct link between RT and SSD within the same trial means the increase in RT on Partial Stop trials is likely to be directly caused by the later stop signal presentation (i.e. SSD) on these trials, rather than from a general proactive slowing strategy as discussed for the SST. Such a proactive strategy would have also delayed Go RTs in the ARIT, and as discussed above, this was not observed. Overall, SSRT may not be the most appropriate term to describe what is being measured during partial RI as it is capturing more than a simple ‘stop signal reaction time.’

The response to a partial stop cue can improve across sessions. The improvement (reflected in SSRT, SSD and accuracy) indicates participants became better at fulfilling the overall demands on Partial Stop trials. This may be because some, or all, of the sequential process captured by the SSRT (stop, uncouple, reprogram, then go) occurred over a shorter time scale in session two. The overall time required for this process can be reduced on Partial Stop trials of both the ARIT (Wadsley et al., 2019) and SST (Xu et al., 2014) through manipulations to overall task design. Wadsley and colleagues (2019) increased the asynchrony between left and right-side components of the default response, thereby reducing the amount of time needed for response uncoupling during partial RI and reducing the stopping interference effect. Xu and colleagues (2015) also decreased the interference effect by targeting the Go process in partial RI, with specific training to shorten reaction times, although this came at a cost of incredibly short SSDs (averaging 98ms) which likely fundamentally changed partial RI behaviour. In the current study, we saw the improvement without alterations to task design, suggesting multiple sessions alone might be sufficient to increase the efficiency of partial RI processes in young healthy adults. If this working hypothesis is correct, one would expect it to be mirrored by an improvement in neural activity within the various networks activated during partial RI. During complete RI, a proactive shift in the pattern of RI network activation is possible. Regions like the right IFG which are initially recruited during the implementation of RI, can be recruited earlier by inhibition cues following multiple sessions, therefore improving SSRT (Berkman et al., 2014). An increase in GABA mediated short-interval intracortical inhibition in the primary motor cortex (M1) could also contribute to these improvements (Chowdhury et al., 2020). On the other hand, it is possible participants may have simply become more comfortable with the increased cognitive challenge for partial RI in the current study, thereby improving performance. The inclusion of neuroimaging in future multi-session experimental designs could help distinguish between these possible mechanisms of effect. Nevertheless, our findings indicate that partial RI measures are susceptible to within-individual changes across multiple sessions. This has implications for future study designs that necessitate collecting behavioural measures over multiple sessions.

The extent of any within-individual changes to SSRT across sessions is also potentially relevant in a clinical context. SSRT is sensitive to cortical and basal ganglia impairments resulting from healthy aging (Coxon et al., 2012; Bloemendaal et al., 2016; Coxon et al., 2016) and a wide range of pathologies such as PD (Gauggel et al., 2004; Obeso et al., 2011; Rahman et al., 2021), schizophrenia (Hughes et al., 2012), ADHD (Lipszyc and Schachar, 2010; Senderecka et al., 2012) and OCD (Lipszyc and Schachar, 2010; McLaughlin et al., 2016). It has been suggested that SSRT is a biomarker for specific pathologies and may hold promise for early diagnosis of cortical/basal ganglia dysfunction (McLaughlin et al., 2016; Rahman et al., 2021). However, to identify any impairments in inhibitory control over time because of pathology, natural trends in the SSRT measure over time need to be quantified first in healthy populations. Likewise, any improvements to SSRT as a result of practice need to be identified to quantify additional improvements in inhibitory control as a result of treatment interventions. To this end, the current study examined subtle changes in SSRT over two laboratory sessions, as might commonly be done pre- and postintervention or when using active non-invasive brain stimulation against sham over two sessions. To further understand if the SSRT as measured on these behavioural tasks has potential to detect pathology or effects of treatment in the future, more studies of this kind must initially take place in healthy subjects whilst ensuring the constraining of the Go response.

It is important to acknowledge the presence of some very low SSD values for partial RI trials in the ARIT for our cohort. Whilst these trials are known to be challenging, SSDs of 50 - 200ms point to particular difficulty meeting trial demands, which is somewhat surprising for young healthy adults. Perhaps these participants required a greater number of Partial Stop trials to arrive at their maximal RI efficiency. However, for some participants these low SSDs persisted until the end of session two. It is therefore possible that these values reflect motor or cognitive impairments on these trials. The impairment could be linked with overall trait impulsivity (Aichert et al., 2012) or even indicate underlying subtle but complex deficits in not only inhibitory control but also conflict monitoring and working memory, as can manifest in pathologies such as ADHD (Rapport et al., 2008; Senderecka et al., 2012).

The current study investigated the consistency of SSRT in the ARIT across two sessions. During complete RI, the maximal efficiency of the inhibitory process remained unchanged for individuals. Results of the SST highlighted that SSRT for complete RI can be affected by more than purely inhibitory network integrity when Go trial reaction times are not constrained in task design. Partial RI measures were susceptible to within-individual changes across multiple sessions and subsequent studies are needed to explore whether the improvements are driven by changes to neural activity within the underlying networks.

Future research should continue to investigate any within-individual changes to SSRT on the ARIT over a greater number of experimental sessions.

## Author’s Roles

AH and HM participated in all aspects of the development of the research project, statistical analyses and the writing and revisions of the manuscript. NJ participated in conception and design of the study and the review and critique of the final manuscript.

## Statements and Disclosures

The authors declare no competing financial interests and no conflicts of interest.

## Funding sources for study

This study was funded by The University of Birmingham.

## Ethical approval and informed written consent

Ethical approval was acquired from The University of Birmingham Ethics Committee to complete the current study.

## Notes

### Competing Interest Statement

The authors have declared no competing interest.

